# Anti-tumour efficacy of *Musa paradisiaca* flower extracts on DMBA induced Mammary carcinogenesis in female Wistar rats

**DOI:** 10.1101/2022.08.27.502223

**Authors:** Kousalya Lavudi, Hemalatha S, Rekha Rani Kokkanti, Harika G.V. S, Srinivas Patnaik, Josthna Penchalaneni

## Abstract

**Background:** Several reports have shown the beneficial advantages of *Musa paradisiaca* in wound healing activities and other ailments. Previously, our *in vitro* studies validated the anti-cancer activities of Musa flower extracts and confirmed the potential. This thread has led to our current investigation to explore the anticancer potential *in vivo.*

**Purpose:** This study focused on the efficacy of banana florets on DMBA induced breast cancer in female wistar rats.

**Methods:** Induction of tumour using Dimethylbenzanthracene (DMBA) in female wistar rats. Hormonal, antioxidant and anticarcinogenic studies were performed *in vivo.*

**Results:** In our current study, we investigated that tumour induction has an effect in disrupting the estrous cycle in mice which may result by hormonal fluctuation levels. Antioxidant and hormonal analysis *in vivo* revealed the imbalance in estrogen and progesterone levels in untreated group compared to the treated ones. Floral crude extract treatment in vivo has shrunken the tumour volume in flower extract treatment group as well as in standard drug compared to the control. Histopathological staining confirms the disruption of epithelial tissues in tumour induced groups compared to the treated ones. Moreover, Musa floral treatment has shown to revert the damaged tissue morphology in the treated groups compared to the saline treated one. In-vitro studies in MCF-7 and MDA-MB-231 breast cancer cell lines has shown the potent anti-tumorigenic activity using Musa floral extracts.

**Conclusion:** Taken together, our finding confirmed that banana flower extracts showcase anti-carcinogenic activity against breast cancer both *in vitro* and *in vivo.* Tumour induction in mice has an effect in showing the disruption of estrous cycles.

**Highlights:**

- *Musa paradisiaca* crude ethanolic flower extracts have a potential in reducing the tumour growth.
- DMBA induced cancer has a deleterious effect on estrogen cycles in female wistar rats.
- Disruption of epithelial morphology was observed in tumour induced wistar rats.
- Treatment with crude Musa flower extracts on cancer induced rats shows the reduced level of damage and oxidative stress compared to the no treatment group.

## INTRODUCTION

Breast cancer is considered as a top-ranking cancer in women world-wide accounting nearly 23% of total cancer cases (Comsa et al., 2015). Several treatments have been carried out to keep this disease in check, yet however the rate of incidence is high in most of the women over 45 years of age. Primary treatment strategies include surgery, chemotherapy, and targeted therapy (McDonald et al., 2016). Several FDA approved medications are in use as a front-line treatment. However, ethnomedicine usage has its vast advantages in treating several ailments relative to standard drugs, which might not only protect the surrounding healthy tissue environment, but also shows fever side effects. Current therapeutic options are ineffective, hence there is a high demand for research in this area.

*Musa paradisiaca,* generally known as banana, is a well-known tropical herbaceous plant of the Musaceae family (Probojati et al., 2021) and a major source of food all over the world (Yusoff, 2008). The banana is considered as a dessert cultivar, but the plantain is known as a culinary cultivar (Oyeyinka and Afolayan, 2020; Vu et al., 2018). It is farmed largely for its fruit and natural fibres and is widely spread around the world (Khoozani et al., 2019). *Musa paradisiaca* have been using in traditional medicine where almost all parts of the plants have medicinal properties. (Mondal et.al 2021; Chinatmunnee et.al 2012; Ghorbani et.al 2012). Banana plant is used as an anti-ulcerogenic, anti-microbial, anti-urolithiatic activity, analgesic, hypoglycaemic and exhibits wound healing properties and it has been linked to a lower incidence of colorectal cancer, renal cell carcinoma, and breast cancer in women (Mathew et al, 2017). It offers a lot of promise when it comes to lowering the risk of hypertension, hyperlipidaemia, stroke, and restoring normal bowel activity, protection against Alzheimer’s disease, kidney health, Immunity booster, and weight reduction and reduces bleeding associated with menorrhagia and reduces risk of certain cancers (Bhaskar et al., 2011a). Traditionally it is used in intestinal lesions, uraemia, nephritis, gout, cardiac disease, hypocholesterolaemia activity and green fruit has been shown to have hypoglycaemic properties (Bhaskar et al., 2011b; Corona et al., 2008) due to increased insulin synthesis and glucose utilization.

Cancer is the second biggest cause of illness and mortality worldwide, despite remarkable advances in research and treatment over decades. Previously, many works have been carried on different parts of *M. paradisiaca* plant like antibacterial and toxicity studies from the ethanolic leaf extract (Asuquo and Udobi, 2016, Ariffin et al., 2021), wound healing properties from methanolic stem extract (Amutha and Selvakumari, 2016). Unripe peels and fruits demonstrating anti-ulcerogenic properties (Ezekwesili et al., 2014), hydroxyanigorufone displayed potential chemo preventive properties for cancer (Jang et al., 2002) have been carried out.

In our study, we have focussed on investigating the anti-carcinogenic activity by using ethanol extracted form of banana flower florets and to validate its further effects on estrous cycle and antioxidant activity. We also reported the anti-tumorigenic efficacy in DMBA induced female wistar rats through H&E staining method. *In vitro* anti-cancer efficacy was performed in both MCF7 and MDA-MB-231 cell lines.

## MATERIALS AND METHODS

### Plant material and extract preparation

The whole inflorescence of *M. paradisiaca* were collected from the local markets in Tirupati, Andhra Pradesh, India. The floral inflorescence has been confirmed by Dr. Nagaraj, Associate professor, Department of Botany, Sri Venkateswara University, Tirupati. Further rinsed thoroughly with distilled water and carefully separated into simple florets. These florets were dried in shade for 2 weeks and ground into coarse powder and stored for further use. The extraction was performed by using soxhlet extractor for 7-8 repeated cycles for 8 h at 50 °C by weighing 30 g of dried powder in 80% ethanol as a solvent. The extracted sample were concentrated by using rotary flash evaporator at 78 °C. The obtained powder was stored in refrigerator (4 degree) for further use.

### Cell culture and maintenance

Human Breast cancer cell lines MDA-MB-231 and MCF-7 were obtained from NCCS-Pune. The cell lines were cultured in Dulbecco’s Modified Eagles medium containing 10% FBS and 1% penicillin and streptomycin and incubated at 37°C in 5% CO_2_ incubator. The medium was replaced at an interval of 48 h. The cells were passaged when 80% of confluency was achieved. All the experiments were performed in between 4-6 passages.

### MTT assay

1*10^5^ cells of MDA-MB-231 and MCF-7 were seeded in 96-well plate and incubated in CO_2_ incubator (5% CO_2_ and 37 °C). After 24 h, the cells were treated with the crude extract at different concentrations. The crude powder extract was dissolved in DMSO and filtered using PVDF syringe filters and added to the wells at different concentrations (5μg-40μg/ ml for MCF-7 cell, and 12.5-200 μg/ml for MDA-MB-231) all the wells were makeup to equal volume with DMEM media and incubated for 24 h. 20 μl of MTT (4,5-dimethylthiazol-2-yl)-2,5-Diphenyltetrazolium bromide) solution (5mg/ml) was added to each well and incubation was performed for 4 h. Then the MTT solution was removed and washed with PBS. The formazan product was collected, and absorbance was measured at 570 nm and O.D was recorded (n= 5). The percentage of cell proliferation was calculated using the following formula:

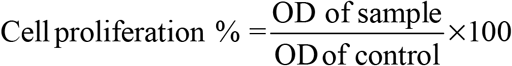

### FACS (Florescence activated cell sorter) analysis

Treated Cells were harvested in 6-well plates at a density of 2 x 10^5^ cells / 2ml into FACS tubes and then incubated. The medium was removed, and 2 ml of culture medium was added and incubated for another 24 h. Later the medium was removed and washed with PBS. 0.25% Trypsin EDTA solution (200 μl) was added and incubated for 3 - 4 min. The cells were harvested directly into 12 x 75 mm polystyrene tubes by adding 2 ml of culture medium and centrifuged at 300 g at 25 °C for 5 min. The cells were fixed in 1ml cold 70% ethanol for 30 min. Then the supernatant was removed, and the pellet was washed twice with the PBS. It was then treated with ribonuclease A (50 μl) for 5 to 10 min at room temperature and cells were analysed by FC in propidium iodide/RNaseA solution.

### Animal study and maintenance

Female wistar rats (n=30) of weight 150-180 g, 8-9weeks old were procured from Sri Venkateshwara Enterprises, Bangalore, India. The animals were housed in plastic cages covered with wire mesh, animal room environment was monitored for temperature 22 ±2 °C, relative humidity of 55 ±5 % and animals were provided with 12 h light and 12 h dark. Animals were divided into 5 groups (n=6) in each group. The body weights were recorded on every alternate day till the end of the experiment. The experimental procedure was approved by Institutional Animal Ethics Committee (IAEC) (Reg. No. 1677/po/Re/S/2012/CPCSEA/33).

Division of groups: (Tumour induction was performed from Group 2-5 using DMBA).

Group 1: No tumour induction, maintained as negative control with saline treatment alone.
Group 2: positive control without any treatment.
Group 3: Treated with low dosage (12.5 μg/ml) of MPFE, *M. paradisiaca* flower crude extract
Group 4: Treated with high dose (20 μg/ml) of MPFE, *M. paradisiaca* flower crude extract
Group 5: Treated with standard drug Tamoxifen (TMA).

Selection of the dose was based on the IC_50_ concentration.one higher dose and one lower dose was selected with reference to IC_50_.

### Vaginal swab smear test

Rats were hold around the thorax and 10 μl of saline solution was injected into the vagina of depth 2-5mm.Vaginal discharge was collected and observed under phase contrast microscope at 100X magnification.

### Hormone analysis or Hormonal assays

Blood samples were collected from the control and treated groups, and thirty minutes after collection, the samples were then centrifuged at 2500 rpm for 15 min at 25 °C, followed by serum collection and stored at −80 °C until analysis. Follicle-stimulating hormone (FSH), luteinizing hormone (LH), progesterone, and estrogen were measured in a clinical laboratory, Tirupati, Andhra Pradesh using the Automated Chemiluminescence Immunoassay method (Hospivision Fully Automatic Immunoassay Analyzer CLIA, A-6, Delhi, India) according to the manufacturer’s instructions.

### Haematology study

#### Total erythrocyte count

Total erythrocyte count was determined by hemocytometry following the method of Gottfried and Gerard (1987).

#### Total leukocyte count

The blood specimen was diluted in ratios 1:20 with WBC diluting fluid and the cells were counted under haemocytometer at 10X magnification (Gottfried and Gerard, 1987) and reported as the number of white blood cells/mm^3^.

#### Platelet count

The platelet count was determined using the Rees and Ecker (1923) method.

#### Differential WBC Count

A thin smear of blood specimen was prepared and stained with Leishman’s staining solution for 2-3 min (Gottfried and Gerard, 1987

#### Estimation of haemoglobin

The estimation of haemoglobin was done using Sahli’s acid hematin method with the help of Hb comparator box and results were expressed in grams per decilitre (g/dl).

#### Packed cell volume (PCV)

The ratio of red blood cells to the whole blood volume was estimated using calibrated micro haematocrit centrifuge at 10,000 rpm for 3 to 5 min. The tube was removed immediately and recorded the PCV to avoid re-dispersion of cells.

### Antioxidant study

We have performed Lipid oxidase peroxidation (LPO), super oxide dismutase assay (SOD), Catalase (CAT), SGOT, SGPT, GPX, ALT and AST as mentioned in the references without any modifications. LPO assay-Niehius and Samuelson (1968), SOD assay, Beauchamp and Fridovich (1971). CAT assay was performed as per the protocol described by Sinha (1972 SGOT activity was determined according to Reitman and Frankel (1957). SGPT activity was performed using the protocol described by Kumar et al (1999), GPx was performed based on the method mentioned by Paglia and Valentine (1967). AST and ALT as mentioned by Arslan et al. (2009).

### Histopathological Studies

A portion of mammary gland tissue of the control, DMBA control (tumour) and treated groups of rats with Tamoxifen and crude extract of *M. paradisiaca* were stored in containers for 12 h in 10% formalin solution and subjected to histopathological studies. Initially the materials were fixed in 10% buffered neutral formalin for 48 hours and then with bovine solution for 6 hours. Paraffin sections were taken at 5 mm thickness processed in alcohol xylene series and was stained with alum haematoxylin and eosin. The slides were observed microscopically for histopathological studies.

## RESULTS

### Anti-tumour activity of *Musa paradisiaca* flower extracts on breast cancer cell lines

MCF-7 and MDA-MB-231 cell lines when treated with MPFE has shown significant cell death. This confirms the anti-proliferative activity of the plant extract against breast cancer cell lines (Fig. 1A). Our results have shown the similar kinds of activity with other reports against various cancers (Kim et al., 2022). The IC_50_ values for MDA-MB-231 and MCF-7 are 90.32 μg/ml and 39.39 μg/ml. These results have shown decrease in the percentage of cell viability with the increase in the concentration of the crude extract (Fig. 1B). This data confirmed that crude extracts have potential cytotoxic properties on the cell lines and inhibit the growth of the cells at higher concentration by arresting the cell growth.

**Figure 1A:**
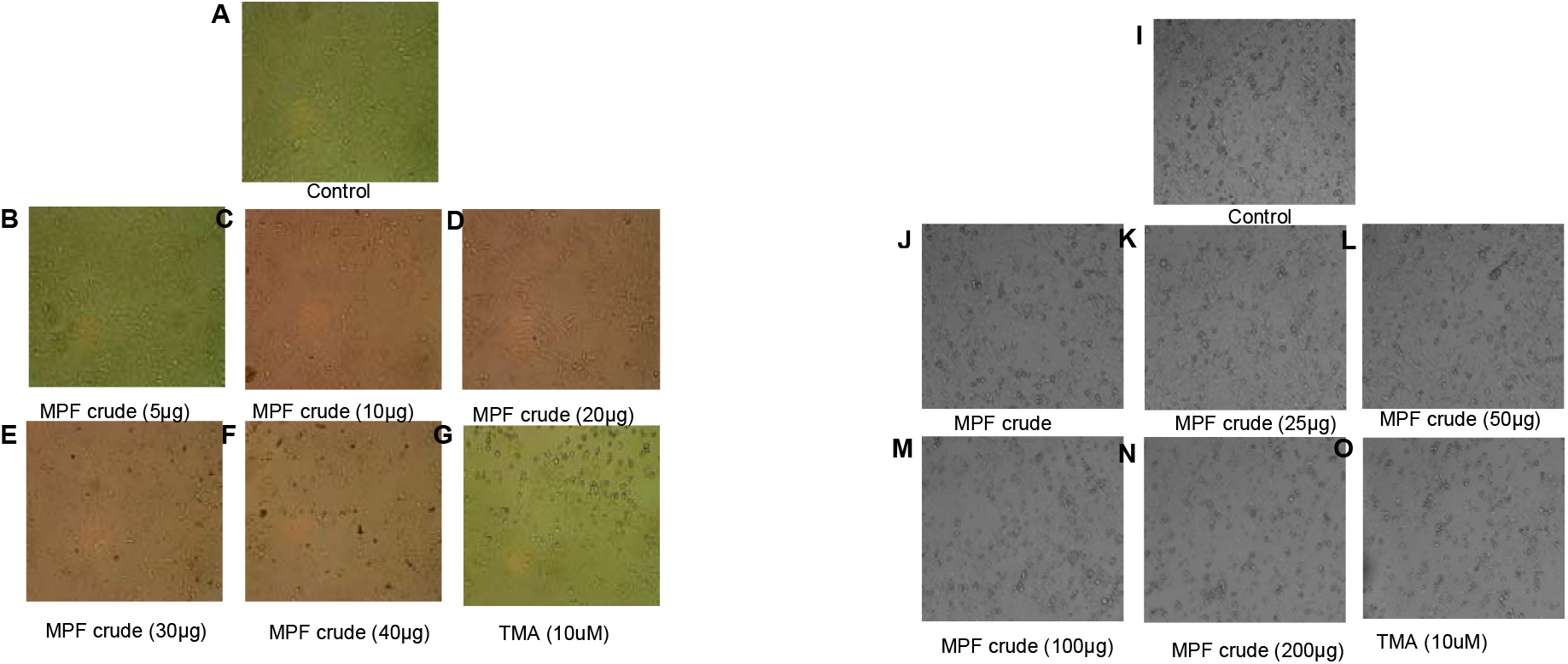
shows the decrease in cell proliferation capacity after treating with respective concentrations of MPF extracts (A-G) MCF cell lines, (I-O) MDA-MB-231 cell

**Figure 1B:**
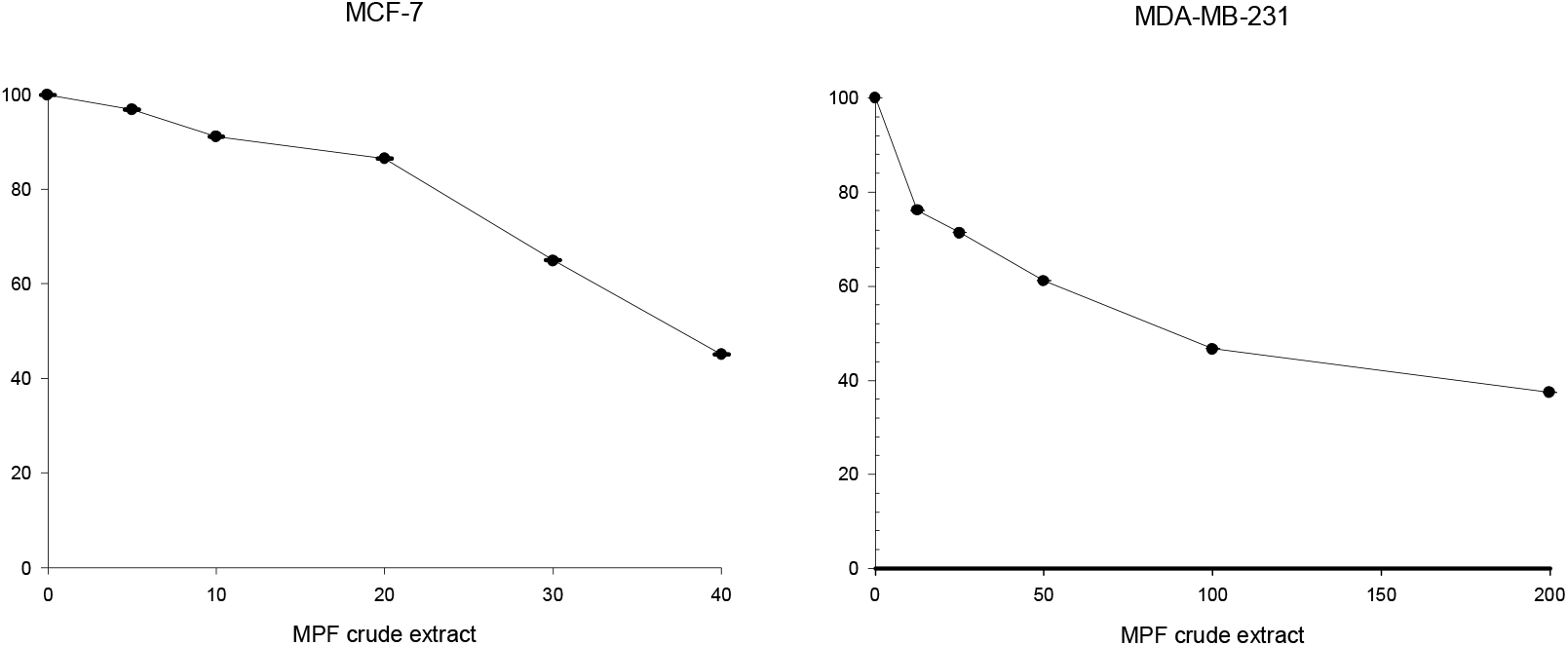
shows the cell sensitivity cure in both MCF and MDA-MB-231 cell lines against MPF extracts.

### Flow cytometry

Flow cytometric analysis was performed for MCF-7 cell line only. Most of the cells were arrested at G1/G2 phase. Untreated cells were arrested at G0/G1 and G2/M phases. This is clear evidence that *M. paradisiaca* flower extract is showing a potential anti-proliferative activity against MCF-7 cell lines (Fig. 2).

**Figure 2.**
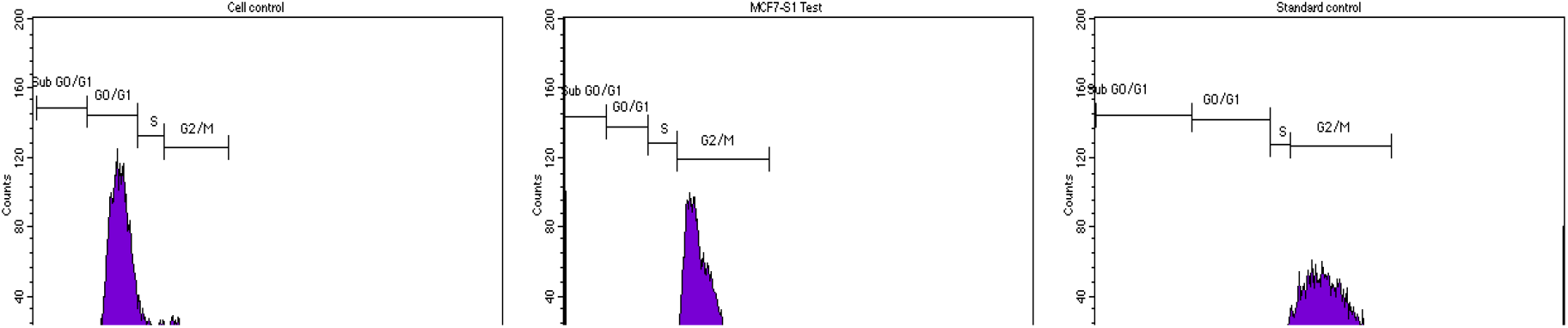
represents the FACS activity in MCF-7 cells treated with MPF extracts.

### Vaginal Swab Smear Test

Vaginal swab smear test was performed to study oestrous cycle in female rats (Fig. 3). The test was performed consequently for 4 days with reference to the photo periodic cycle. The test was conducted before and after inducing the rats with DMBA and all the 4 phases were observed. The results have shown that the cycle was disturbed in the rats induced with DMBA which clearly shows that DMBA affects the hormonal imbalance.

**Figure 3:**
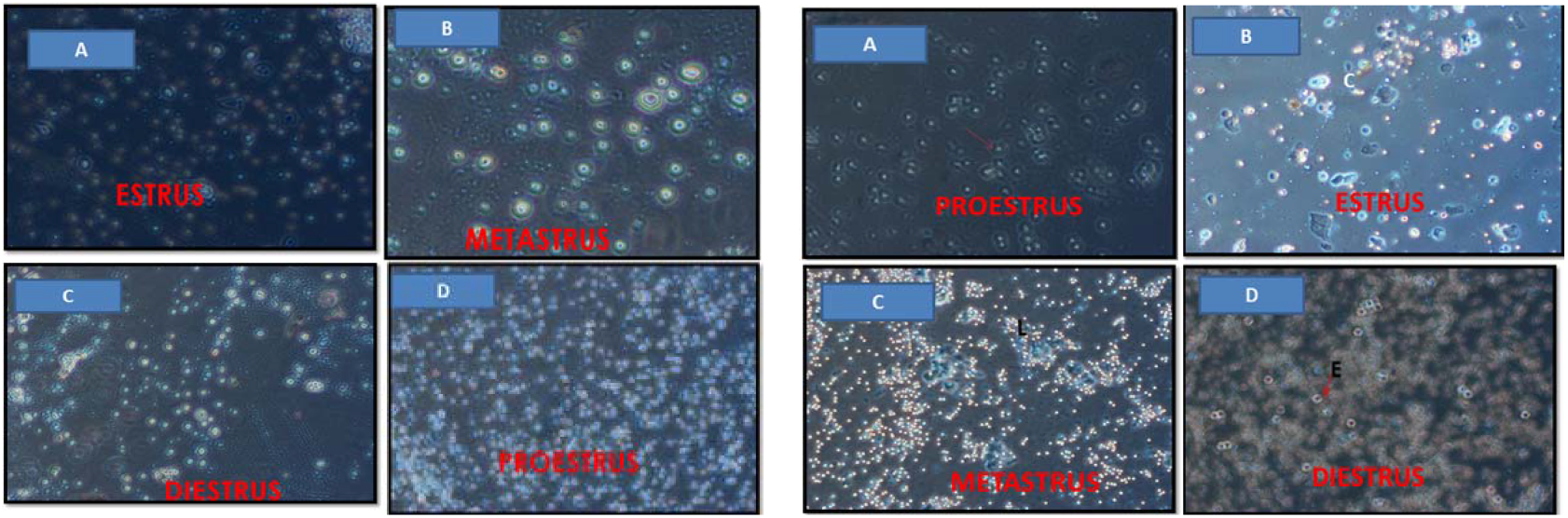
Represents the oestrous cycle in female wistar rats A-Before tumour induction and B-After tumor induction.

### Body weights of rats

Administration of lower and higher dose of MPFE extract, results no mortalities or evidence of adverse effects implying that the crude extract was nontoxic. Throughout the treatment phase, no changes were observed in the behaviour pattern or clinical sign (Table 1). In group 1 and 2, slight increase in the body weight was observed, but in group 3, 4, and 5 decrease in the body weight was observed. Reduction in the body weight is possible due to the development of cancer induced by DMBA. The results suggest the protective role of MPFE with no significant adverse effects associated with treatment.

**Table 1:** Body weights of rats before and after treatment. Body weights of rats. The weights of the rats were represented in grams. The data was analysed, and average weights (n=6) were given with standard errors “±”.

| Body Weights (gm) | Group 1 | Group 2 | Group 3 | Group 4 | Group 5 |
| --- | --- | --- | --- | --- | --- |
| Before Treatment | 179±11.143 | 268.3± 10.3 | 263.5± 17.93 | 280.3 ±12.41 | 279.16 ± 15.27 |
| After Treatment | 182.5±9.35 | 276.6±12.11 | 236.6± 14.71 | 232.5±15.411 | 223.3 ±13.29 |

Body weights and tumour volumes were measured for all groups (Table 2). Further rats were sacrificed at the end point and tumour effecting the mammary glands were isolated. These results are evident that in group 2, tumour volume is high as expected because of no treatment. However, in groups 3 and 4, which received different concentrations of MPFE shown decreased tumour volumes. This confirms the anti-cancer efficacy and tumour volume reduction capacity of MPFE in vivo.

**Table 2:** Effect of treatment on tumour weights.

| S. No | Groups | Tumours(g) | Centimetres (cm) |
| --- | --- | --- | --- |
| 1. | Group 1 | – | – |
| 2. | Group 2 | 6.35 | 1.3 |
| 3. | Group 3 | 4.24 | 1.0 |
| 4. | Group 4 | 0.92 | 0.5 |
| 5. | Group 5 | 0.20 | 0.1 |

#### Effect of *M. paradisiaca* extract on antioxidant assays

LPO are oxidation products of phospholipids and poly unsaturated fatty acids play an important role in physiological processes like antioxidants, immune regulators, and anticancer factors. GPx is an antioxidant enzyme class with the capacity to scavenge free radicals which are involved in the termination reaction of ROS pathway. SOD and CAT enzyme activities were checked as it plays an important role in relieving oxidative stress.

The results (Figure 4) have shown that group 1 have the highest activity in all the enzyme assays followed by group 5 and 4. Groups 3 and 4 which were treated with MPFE have shown the better enzyme activity when compared with DMBA treatment alone. Group 2, which has received lower concentration of MPFE has shown overall reduced enzymatic activity. Group 4 mice have shown higher efficacy in preventing the DMBA deleterious effects compared to others. The results have indicating that flower extract of *M. paradisiaca* on treatment enhances the enzyme activity to act against the free radicals.

**Figure 4:**
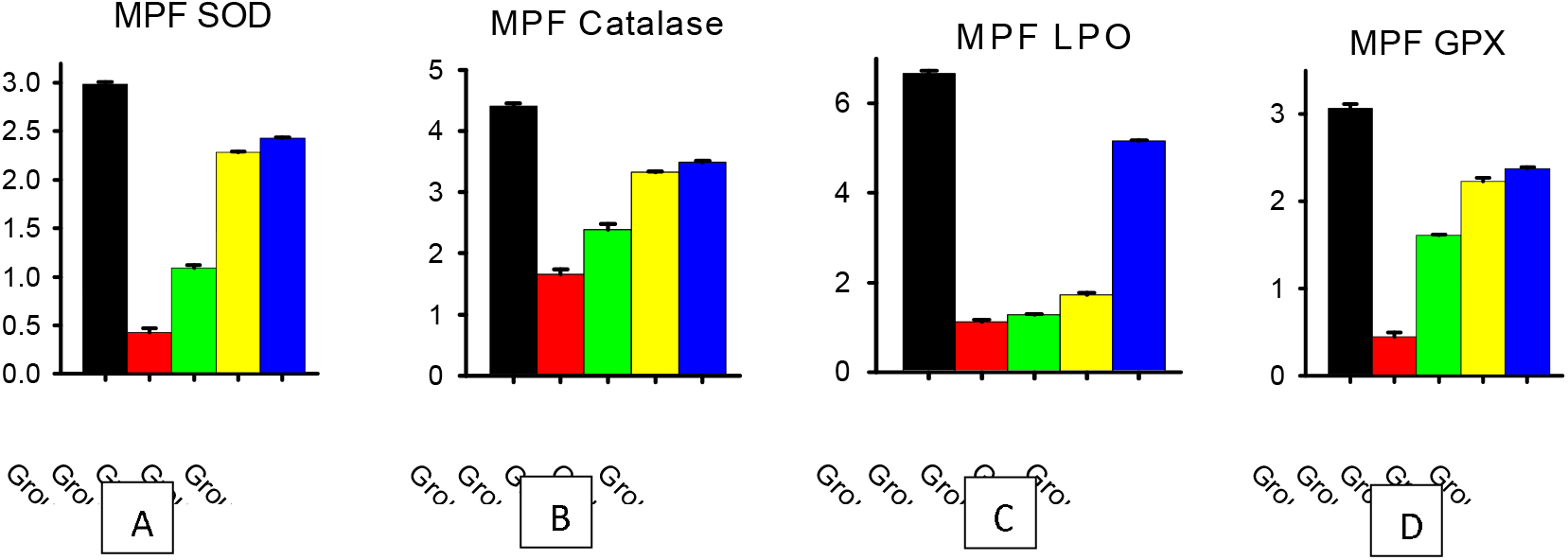
Enzyme activity of blood serum of rats (A, B, C and D).

#### Effect of *Musa paradisiaca* flower extract on AST and ALT activity

The activity of AST and ALT of control and treatment groups were analysed (Figure 5). The results have shown that group 2 has higher activity when compared to others in both the assays. Difference was insignificant in the treated groups (3-5).

**Figure 5:**
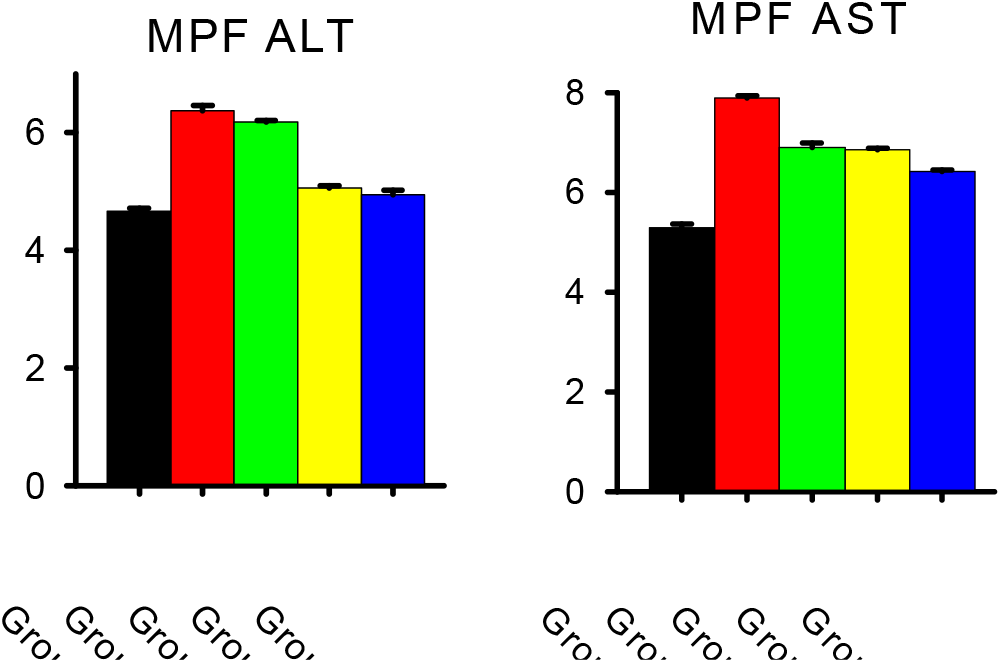
AST and ALT activity of *M. paradisiaca.*

#### Hormone levels estimation

The hormones levels were checked for the control and treatment groups (Table 3). The results have shown that FSH and LH have shown similar hormone levels with slight changes. DMBA treatment has shown higher values than the reference values.

**Table 3:** Effect of *M. paradisaica* on hormones level. Determines the hormones levels were checked for control and treated groups. The average values were given.

| Hormones | Group 1 | Group2 | Group3 | Group4 | Group5 | Reference ranges |
| --- | --- | --- | --- | --- | --- | --- |
| FSH | 2.3 | 2.9 | 2.4 | 2.5 | 2.5 | 2.00±0.12 |
| LH | 2.4 | 2.7 | 2.65 | 2.49 | 2.45 | 2.46±0.3 |
| Estrogen | 23.35 | 38.9 | 37.5 | 35.4 | 34.9 | 29.7±7.61 |
| Progesterone | 11.37 | 6.91 | 10.35 | 10.56 | 10.59 | 12.20±2.18 |

Estrogen levels were higher in group 2 followed by group 3, 4, 5. Control group have shown less estrogen levels compared to reference values. Progesterone levels were observed differently in the groups. Maximum progesterone levels were higher in control group 1 followed by group 4, 5 and 2. Higher dose of MPFE treatment was more effective as compared to lower dose in preventing DMBA effects signifying the protective role of MPFE.

#### Haematology studies

Comparing the haematology studies with stand reference values, haemoglobin, leucocytes count, neutrophils, monocytes, RBC count, packed cell volume and MCHC were within the reference range. Remaining all the studies have shown difference in the results, were given in table 4. These studies confirmed that metastasis was not observed in fluid connective tissue.

**Table 4:** Haematological analysis: Represents Haematological analysis carried in all the groups. The data was represented for the assays with standard reference values.

| Studies | Group 1 | Group 2 | Group 3 | Group 4 | Group 5 | Reference ranges |
| --- | --- | --- | --- | --- | --- | --- |
| Haemoglobin | 12 | 10.4 | 11.3 | 10.9 | 11 | 10-13gms/dl |
| Leucocyte count | 11400 | 7200 | 8100 | 8600 | 7600 | 10000-12800 cells/cumm |
| Neutrophils | 42 | 40 | 23 | 39 | 50 | 23- 44.9% |
| Lymphocytes | 56.1 | 93 | 81 | k56 | 79 | 56.1 – 77% |
| Eosinophils | 01 | 00 | 03 | 04 | 04 | 0 -1% |
| Basophils | 00 | 00 | 00 | 00 | 00 | 0 -1% |
| Monocytes | 01 | 01 | 02 | 01 | 01 | 0 -1% |
| RBC count | 9.8 | 07 | 11 | 10.5 | 10.9 | 8-10.4million cells/cumm |
| Packed cell volume | 38 | 21 | 32 | 32 | 31 | 33-50% |
| Platelet count | 520 | 900 | 700 | 850 | 880 | 480-725 lakh cells/cumm |
| MCV | 60 | 76 | 67 | 76 | 79 | 46-65 f l |
| MCH | 16 | 23 | 20 | 21 | 22 | 14-18.5 pg |
| MCHC | 31 | 30 | 30 | 27 | 28 | 28.2-32.4 gm/wt |

#### Histopathological Studies

Histopathological examinations of mammary glands reveal that disruption of connective tissue occurred when compared with the control (Fig.6). Standard drug treated mammary tissues exhibit invasive ductal carcinoma (IDC) which is a benign form of tumour. Untreated mammary tumour samples exhibit a highly disorganized structure and deformed connective tissue. Stromal spaces are clearly visible which confirmed the proliferation rate and metastasis to pancreatic region.

**Figure 6A:**
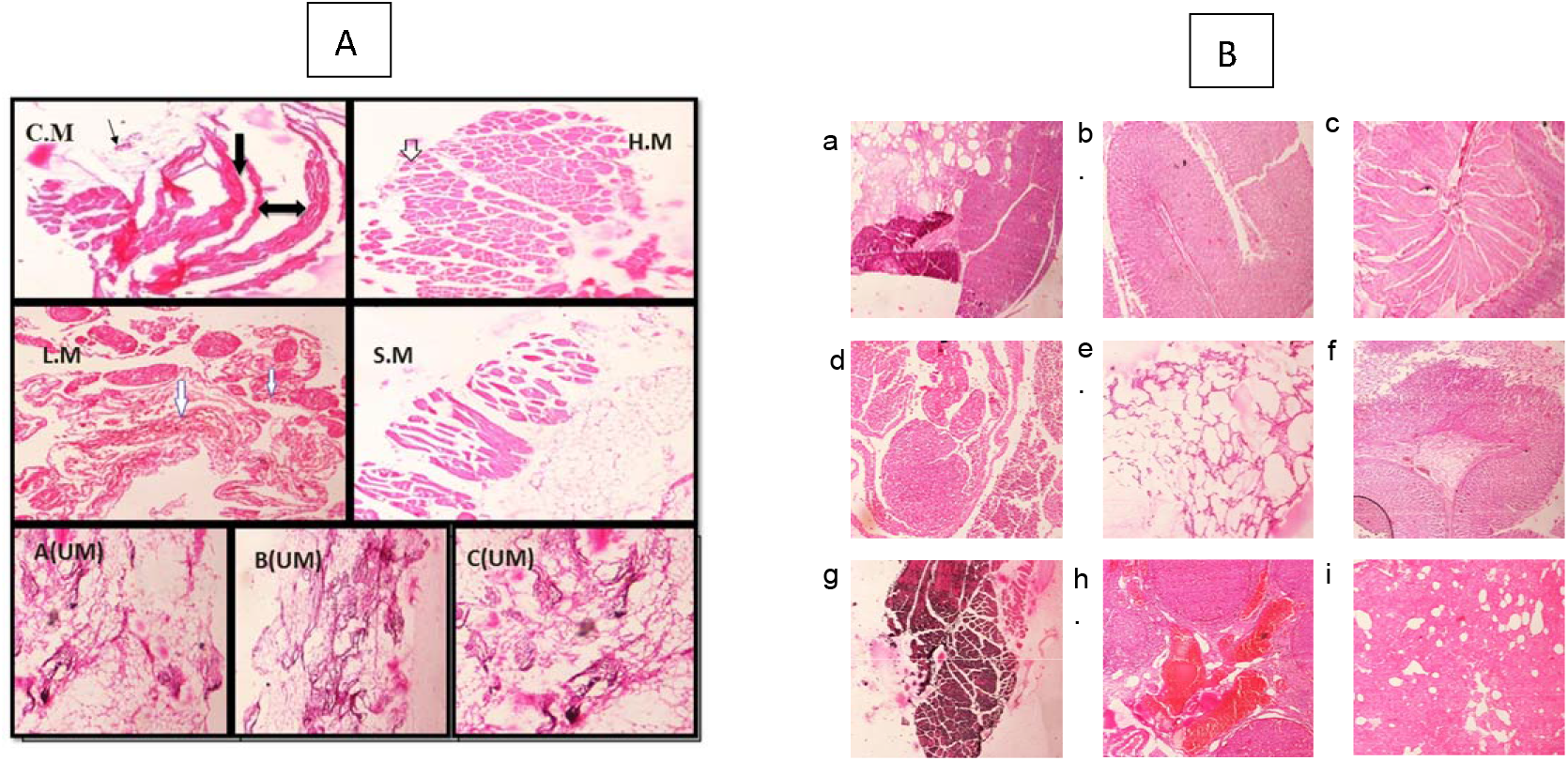
**C.M**-control mammary gland; **H.M**-High dose treated mammary gland; **L.M** – Low dose treated mammary gland; **S.M** – Standard dose treated mammary gland; **A, B and C (UM)** - Untreated mammary gland.

Low dose drug treated tumour samples showed proliferation in few regions whereas high dose treated tumour samples examined the reformation of connective tissue and the proliferation of cells were reduced. Standard drug has also showed the reformation, yet the tumour growth was controlled in beginning stage itself.

## DISCUSSION

There is a large amount of experimental evidence which suggests that consumption of fruits and vegetables lower the risk of cancer. In the present study, DMBA induced mammary cancer in female wistar albino rats were treated with the ethanolic crude flower extracts of *M. paradisiaca* at different concentrations. Cell studies have revealed that the crude extract exhibits good cytotoxic potential against MDA-MB-231 and MCF-7 cell lines and prove as a potent anti-cancer agent. Previously it was reported that the banana peel exhibited significant antitumour activity against the MCF-7 cell lines (Durgadevi et al., 2019), *M. cavendish* green peel hydroalcoholic extract exhibited antiproliferative activity against the MCF-7 cell line (Barroso et al., 2019), hexane extract of *M. sapientum* peel and pulp was observed to be cytotoxic and blocked the proliferation of MCF-7 cells (Dahham et al., 2015). The blossom extracts were also found to show cytotoxic and antiproliferative activities against several cancer cells such as human colon cancer cells, HT29, and HCT-116, as well as HeLa and breast cancer cells (Arun et al., 2018; Nadumane and Timsina, 2015). The present study also correlates with the previous work results as the crude extract of *M. paradisiaca* act as good anticancer agent but at higher concentration.

The decrease in the body weight in rats was observed in the treated groups compared to control; this might be due to the development of the tumour in the rats. The weight of tumour is found to be less in tamoxifen treatment followed by high concentration crude extract treatment. Our understanding towards this could be because of standard drug might affect the growth of tumour *in vivo.*

Moreover, in this study we tried to evaluate the correlation between antiangiogenic and antioxidant activity of banana extracts. The ethanol extracts of banana flower also exhibited potent antioxidant properties. Antioxidant assays reveal that in untreated group of rats AST and ALT levels were elevated as these serum enzyme concentrations are raised during the cellular damage and oxidative stress. The antioxidant enzymes such as SOD, CAT and GPx were assessed to study the potential role of MPFE against DMBA induced oxidative stress in female wistar rats. The reduction in the levels of enzymes suggests their active role in response to stress compared to control conditions. Decreased levels of LPO in DMBA induced groups is due the increased levels of estrogen in group 3 to 5. According to reports by Dominguez et al., 2005; Das, 2002, elevated estrogen levels inhibit LPO activity in breast cancer cells associated with increase in cell division. The findings of Mandlekar and Kong, 2001; Mandlekar, 2000; Gundimeda, 1996 confirms the interventions to LPO other than hormones and antioxidants also affecting the behaviour of breast cancer cells including the tamoxifen treated groups.

Hormonal analysis showed that the levels of LH, FSH and estrogen are elevated, and progesterone was decreased in group 2 which has not receive any treatment. This may be due to the action of DMBA in changing the hormone production mechanism regulating hypothalamic pituitary axis (HPA) because of oxidative stress in treated groups (Afsar et al., 2017). Our previous study on PA-1 ovarian cancer cells using methanolic extracts of banana peudostem has shown potential anti-cancer properties. (Rohini et.al 2021). Correlation to the previous studies, our current results showed that the crude extracts possess anti-proliferative activity and proliferation inhibition was significantly increased in dose-dependent manner.

## CONCLUSION

The present study has explored the role of extracts of *M. paradisiaca* flower in regulating the cell proliferation *in vitro* and *in vivo.* The histopathological studies showed that the tumour was developed and exhibits a high mitotic rate as well as stromal invasion and affects the pancreatic region of the female wistar untreated rats. However, further studies are necessary to confirm the mechanistic process of invasion and the role of Musa flower extracts in apoptosis and signal transduction mechanisms. Advanced studies are needed to confirm the activity of *M. paradisiaca* in personalized medicine to treat human samples.

## ACKNOWLEDGEMENTS

We would like to acknowledge Department of Biotechnology, New Delhi, India for providing contingency amount.

## CRediT author statement

Conceptualization: Josthna Penchalaneni (PJ), Kousalya Lavudi (KL), Methodology: Hemalatha Salika (HS), Kousalya Lavudi (KL). Investigation: Hemalatha Salika (HS), Kousalya Lavudi (KL). Writing original draft: Rekha Rani Kokkanti, Kousalya Lavudi. Visualization: Harika GVS, Hema Salika. Supervision: Srinivas Patnaik

## Abbreviations

CLIA: Chemiluminescence Immunoassay analyzer
DMBA: Dimethylbenzanthracene
DMEM: Dulbecco’s modified eagle medium
FACS: Florescence activated cell sorter
FBS: Foetal bovine serum
FDA: Food and drug administration
FSH: Follicle-stimulating hormone
H&E: Haematoxylin and eosin
LH: Luteinizing hormone
HPA: Hypothalamic pituitary axis
IAEC: Institutional animal ethics committee
IC50: inhibitory concentration
IDC: Invasive ductal carcinoma
MCHC: The mean corpuscular haemoglobin concentration
MPFE: *Musa Paradisiaca* flower extract
NCCS: National centre for cell Science
PCV: Packed cell volume
PBS: Phosphate buffer saline
PVDF: Polyvinyl difluoride
ROS: Reactive oxygen species
TMA: Tamoxifen

